# Glis3 is a modifier of cyst progression in autosomal dominant polycystic kidney disease

**DOI:** 10.1101/2025.06.08.658051

**Authors:** Zemeng Wei, Jianlei Gu, Xin Tian, Chao Zhang, Hongyu Zhao, Stefan Somlo

## Abstract

**Background:** Autosomal dominant polycystic kidney disease (ADPKD) is caused by mutations affecting polycystin-1 (PC1) or -2 (PC2). The existence of a ‘cilia-dependent cyst activation’ (CDCA) pathway has been identified by demonstrating that structurally intact primary cilia are crucial for cyst growth following loss of polycystins. We previously used translating ribosome affinity purification (TRAP) RNA-Seq on pre-cystic mouse kidneys to determine the translatome that meet the criteria for CDCA and identified *Glis2* as an early effector of polycystin signaling. Here, we investigate the potential role of *Glis3* in ADPKD, which, while not transcriptionally altered, encodes a cilia-localized transcription factor belonging to the same family of genes as *Glis2*.

**Methods:** We used live cell confocal imaging to study the subcellular localization of Glis3 in the presence or absence of *Pkd1*. We used *Glis3* conditional knockout mice to investigate the genetic interaction between *Glis3* and *Pkd1*. Changes in gene expression and chromatin accessibility were assessed by RNA-Seq and ATAC-Seq on an allelic series of Glis3 and Pkd1 kidney selective inactivation models at 7-weeks age to evaluate the function of Glis3.

**Results:** The ciliary localization of Glis3 is not affected by Pkd1 mutation status. Kidney selective *Glis3* by itself does not affect kidney structure or function, but dual inactivation of *Glis3* and *Pkd1* significantly worsen polycystic kidney disease. Integration of transcriptomic profiling and chromatin accessibility assays suggested that kidney tubule specific Glis3 inactivation results in dysregulated fatty acid metabolism and circadian.

**Conclusions:** Glis3 is a primary cilium localized transcription factor that genetically interacts with *Pkd1* and modifies kidney epithelial cell metabolism and circadian function.

## Introduction

Autosomal dominant polycystic kidney disease (ADPKD) is one of the most common human monogenetic diseases, affecting over 12 million people worldwide. Most cases of ADPKD result from mutations in either *PKD1* [^1,2^] or *PKD2* [^3^], respectively encoding the proteins polycystin-1 (PC1) and polycystin-2 (PC2). ADPKD is mainly characterized by cysts originating from the epithelia of kidney tubules, and is varyingly associated with extrarenal manifestations including bile duct cysts and intracranial aneurysms^4^. The precise functions of polycystins and the precise molecular mechanisms underlying ADPKD remain incompletely understood. A number of known pathways have been implicated in ADPKD, including canonical Wnt signaling, mammalian target of rapamycin (mTOR), cyclic AMP-dependent signaling, calcium signaling, G-protein signaling, and disordered cellular metabolism^5–7^. There is general consensus that polycystins expressed on primary cilia are an essential component of the disparate molecular signaling events associated with cyst formation. Genetic studies in mice have shown that structurally intact cilia are a critical requirement for relatively rapid cyst growth following the loss of polycystins^8^. This has led to the inference of a “cilia dependent cyst activation” (CDCA) pathway^8^ that acts as an effective gain of function signal downstream of polycystin inactivation in the setting of intact cilia. We previously performed translating ribosome affinity purification (TRAP) RNA sequencing on pre-cystic mouse kidneys to define a translatome that is associated with CDCA pattern of expression. From this, we identified Glis2 as an early effector of polycystin signaling^9^.

Glis2 belongs to the Gli-similar (GLIS) Kruppel-like zinc finger transcription factor family^10^ which has two other members, Glis1 [^11^] and Glis3 [^12^]. Glis1-3 share structural homology in the five tandem Cys2-His2 zinc finger motifs with the Gli transcription factors in Hedgehog signaling, with the rest of protein sequences being divergent^10–12^. Glis3 is abundantly expressed in the kidney, while Glis1 is expressed at a relative low level^13^. Recessive mutations in Glis3 are associated with a rare syndrome of neonatal diabetes and congenital hypothyroidism (NDH)^14^, in which some patients develop cystic kidneys^15^. Two Glis3 null mice were generated independently, in which either the fifth zinc finger motif^16^ or all five zinc finger motifs^17^ are deleted. Both of the homozygous Glis3 null mice are born in correct Mendelian ratios but die shortly after birth because of extra-renal phenotypes although they also exhibit cyst formation in renal tubules and the glomerulus^16,17^. More importantly, Glis3 has been localized in both the nucleus^18^ and the primary cilia^16^. Given the coordinated functioning of Gli1-3 in Hedgehog signaling and the importance of Glis2 as a CDCA effector in ADPKD, we sought to determine whether Glis3, while not transcriptionally altered in ADPKD, is nonetheless a cilia-localized transcription factor that is part of the ADPKD signaling pathways.

## Methods

### Mouse strains and procedures

All experiments were conducted in strict accordance with the guidelines and procedures set by the Institutional Animal Care and Use Committee (IACUC) at Yale University. The mice used are >95% congenic on the C57BL/6J background. Mice of both sexes were used. The following strains of mice used in this study were previously described: *Pkd1^fl^*[^19^], *Pax8^rtTA^* (JAX Strain # 007176) and *TetO^Cre^* (JAX Strain # 006234). *Glis3^fl^* mice were produced for this study by introducing flanking *lox*P sites around exon 6 using CRISPR/Cas9 gene targeting. Founders were identified by PCR genotyping and verified by sequencing of PCR products. Gene inactivation in the early-onset models with the *Pax8^rtTA^; TetO^Cre^* digenic system was induced with 2 mg/ml of doxycycline in drinking water supplemented with 3% sucrose provided to the nursing dams for two weeks from postnatal day 0 (P0) to P14. All pups were examined at P14. Gene inactivation in the adult-onset models of *Pax8^rtTA^; TetO^Cre^* was induced with 2 mg/ml of doxycycline in drinking water supplemented with 3% sucrose given to mice for two weeks from P28 to P42. Adult-onset model mice were euthanized at 7, 14, 18, and 24 weeks of age. Genotyping was done on DNA isolated from toe clips. Copy number of Pax8^rtTA^ and TetO^Cre^ was determined as previously reported^9^. Blood samples were obtained by ventricular puncture. Serum was separated using Plasma Separator Tubes with lithium heparin (BD Biosciences, Cat. no. 365985). Serum urea nitrogen was analyzed by the Animal Physiology Core of the Department of Internal Medicine at Yale University. One kidney was snap-frozen for protein and mRNA extraction, and the other kidney was fixed in 4% paraformaldehyde for histological analysis. Sagittal sections of kidneys were processed for hematoxylin and eosin (H&E) staining. Cystic index was calculated as previously described^20^.

### Primary cell culture

Primary cells from kidneys were isolated and cultured as previously described^9^. Cells were let attach for 48-72 hours (h) and then treated with doxycycline (Sigma, Cat. no. D9891-100G) 1 µg/ml in the culture media for 72 h. Cells were serum starved to promote cilia formation in 0.1% FBS containing media for 24 h prior to preparation of protein or RNA.

### cDNA constructs, transfection, and cell culture

The full-length cDNA of mouse Glis3 (NCBI Accession NM_001404124.1) was cloned into the pLenti-CMV-GFP-Blast (659-1) vector (Addgene, Cat. no. 17445). Wild type and Pkd1^KO^ IMCD3 cells stably expressing the Nphp3^(1–200)^-mApple cilia marker^9^ were transfected with the Glis3 construct through electroporation and were selected with G418 (500 µg/ml, Sigma-Aldrich, Cat. no. G8168-100ML) 48 h after electroporation. All cell lines were cultured in DMEM high glucose (Thermo Fisher Scientific, Cat. no. 11965092) with 5% FBS and were serum starved in 0.1% FBS containing media for 24 h prior experimentation.

### Immunocytochemistry

WT and Pkd1^KO^ IMCD3 cells were imaged as non-fixed live cells. Immunofluorescence of kidney cryosections (5-7 µm) was performed according to standard procedures. Briefly, slides were permeabilized with 0.2% Triton X-100 in PBS for 30 min and blocked with 0.1% BSA and 10% goat serum in PBS for 1 h at room temperature, followed by primary antibody incubation overnight at 4 °C and secondary antibody incubation for 1 h at room temperature. All images were acquired on Nikon Eclipse Ti (Nikon Instruments Inc, Japan) equipped with Yokogawa CSU-W1 spinning disc and Andor solid state lasers (Andor Technology, UK), using the NIS-Elements AR software (Nikon, Version 4.30.02). The following antibodies and lectins were used: Alexa Fluor^®^647-conjugated (Santa Cruz Biotechnology, Cat. no. sc-23950 AF647) acetylated α-Tubulin (1:1000); rhodamine *Dolichos biflorus agglutinin* (DBA) (1:200, Vectors Laboratories, Cat. no. RL-1032); and FITC *Lotus tetragonolobus agglutinin* (LTA) (1:200, Vectors Laboratories, Cat. no. FL-1321). Hoechst 33342 (1:5000, Molecular Probes, Cat. no. H3570) was used for nuclei staining.

### Protein preparation, immunoblotting, and immunoprecipitation

Fractionation of cells into cytosolic and nuclear fractions were done using the NE-PER^TM^ Nuclear and Cytoplasmic Extraction Reagents (Thermo, Cat. no. 78833) with manufacturer’s instructions. Protein concentrations were measured with Protein Assay Dye Reagent Concentrate (Bio-Rad, Cat. no. 5000006). Equal amounts of total protein were loaded and separated in 4-20% Mini-Protean TGX Precast Gels (Bio-Rad, Cat. no. 4568094) and transferred to Nitrocellulose membrane (Bio-Rad, Cat. no. 1620115). Membranes were blocked with 5% milk for 1 h, incubated with primary antibodies overnight at 4 °C, and were incubated with secondary antibodies for 1 h at room temperature. Membrane stripping and re-probing were done using Restore PLUS Western Blot Stripping Buffer (Thermo Scientific, Cat. no. 46430). The images were acquired with LI-COR Odyssey Fc Imaging system. The following primary antibodies were used: rabbit anti-Glis2 (YNG2 [^9^], 1:20000) and rabbit anti-Lamin A/C (1:1000, Cell Signaling Technology, Cat. no. 2032S). Secondary anti-rabbit horseradish peroxidase (HRP) (1:5000, Jackson ImmunoResearch, Cat. no. 711-035-152) were used.

### RNA isolation, RT-qPCR, and RNA-Seq analysis

Total RNA from cells or whole kidneys were isolated using Trizol (Themo Fisher Scientific, Cat. no. 15596026) and RNeasy Mini Kit (Qiagen, Cat. no. 74104) and used for cDNA synthesis by iScript cDNA synthesis kit (Bio-Red, Cat. no. 1708890) or used for RNA sequencing directly. RT-qPCR was performed using iTaq Universal SYBR green Supermix (Bio-Rad, Cat. no. 18064022) in CFX96 Touch Real-Time PCR detection system (Bio-Rad, USA). The following primers for RT-qPCR were used:

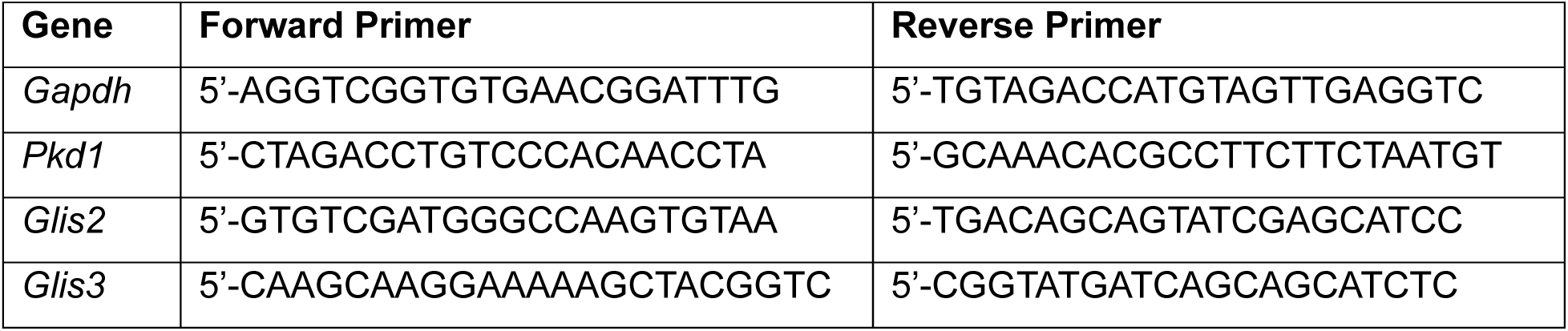

RNA sequencing library was prepared using the ribosomal depletion method (KAPA RNA HyperPrep Kit with RiboErase, Roche). Sequencing was run on the Illumina NovaSeq 6000 platform using 150 bp paired end reads with read depth 100 million reads per sample. RNA sequencing analysis was performed based on previously published pipeline^9^. Briefly, raw fastq files were processed using *fastq* tool^21^. Sequencing reads with low-quality bases were trimmed and filtered. Cleaned reads were aligned using *STAR*^22^ (version 2.7.9) with the mouse reference genome gencode version GRCm38.p6 with vM25 gene annotation. Expression quantification for aligned reads was performed using *featureCounts*^23^ (version 2.0.0). We only focused on protein coding genes. The filtered read counts matrix was normalized by the transcripts per million (TPM) method. Detection of differentially expressed genes was performed using R package *DESeq2* [^24^] with false discovery rate (FDR) ≤0.05 as the significance threshold. Heatmaps were generated using the *pheatmap* package in R with input matrix consisted of log_2_(TPM+1) transformed gene expression values. Gene ontology analyses and gene set enrichment analyses were performed using the *clusterProfiler*^25^ R package.

### ATAC-Seq processing and data analysis

Assay for transposase-accessible chromatin with sequencing (ATAC-Seq)^26^ was performed based on a modification of the ActiveMotif protocol (ActiveMotif, Cat. no. 53150) and the Omni-ATAC method^27,28^. Briefly, kidney tissues were collected from experimental animals, snap frozen in liquid nitrogen, and stored in −80 °C until proceeding for ATAC-Seq. Samples were resected into small pieces in a container with liquid nitrogen to avoid freeze and thaw and transferred to a homogenization tube with three CK28 beads (Bertin Crop, Cat. no. P000911-LYSK0-A) and 500 μl of ice-cold nuclei preparation buffer (10mM Tris-HCl pH7.5, 3mM MgCl_2_, and 10mM NaCl). Homogenization was done three times at 4500 rpm for 10 s each time with the Precellys^®^ evolution homogenizer (Bertin Crop). After homogenization, the nuclei suspension was passed through a 20-micron strainer, washed twice with buffer at 2000 rpm for 1 min, and resuspended in 500 μl buffer. Nuclei were then checked under microscope and counted for 50,000 nuclei per sample. Tagmentation step follows the ActiveMotif protocol with modifications of using half the amount of tagementation mix and incubating at 37 °C for 20 min. The tagemented DNA were purified and amplified following ActiveMotif protocol. The final ATAC-Seq library was analyzed by the Yale Center for Genome Analysis (YCGA) using a TapeStation to assess size distribution and was sequenced at 50 bp paired end reads for 100 million reads per sample.

ATAC-Seq data were processed based on published pipeline on mouse kidney^29^. In brief, raw sequencing reads were trimmed using *cutadapt* and mapped to mm10 using *bowtie2* [^30^]. Duplicates and mitochondrial reads were marked by *Picard* and removed by *Samtools*. Peak calling was performed using *MACS2* [^31^] with peaks falling into blacklist regions (mm10 blacklist v2) removed. One Pkd1KO+Glis3KO sample was an outlier and was removed from downstream analyses. Transcription start site (TSS) enrichment score showing the aggregate distribution of ATAC-Seq peaks relative to TSSs with flanking regions spanning ±2 kb upstream and downstream of TSSs was performed by *ChIPseeker*^32^. Consensus peaks were generated by requiring overlap of *MACS2*-called peaks in at least three biological replicates (minOverlap = 3). Peak annotation was subsequently conducted using *ChIPseeker* in R as well. Differential accessible regions (DARs) were identified using the *Diffbind* R package and FDR ≤ 0.05 was considered as the threshold for significance. *De novo* motif analysis was done by *HOMER*^33^ (version v4.11). Gene ontology analysis was performed by GREAT^34^. Footprinting analysis was performed using TOBIAS^35^.

### Statistics

Quantitative data were analyzed using either one-way analysis of variance (ANOVA) followed by Tukey’s multiple-comparison test or two-tailed, unpaired Student’s *t* test as indicated in figure legends. All data are presented as mean ± s.e.m, and *P* ≤ 0.05 was used as the threshold for statistical significance. GraphPad Prism (10.3.0) software was used to perform statistical analyses.

## Results

We first determined whether the cilia localization of Glis3 is altered by inactivation of *Pkd1*. We stably expressed C-terminal EGFP tagged Glis3 in isogenic wildtype (WT) and *Pkd1* knockout (Pkd1^KO^) IMCD3 cells^36^ that have stable expression of Nphp3^(1–200)^-mApple cilia marker^9^. In WT IMCD3 cells, Glis3 showed the expected localization in primary cilia (Figure 1a). The pattern of expression was not changed in Pkd1^KO^ IMCD3 cells (Figure 1a) indicating that at the level of light microscopy, the cilia localization of Glis3 is not affected by *Pkd1* inactivation in this epithelial cell line. We next sought to determine if there was functional interaction between *Glis3* and *Pkd1*. The neonatal lethality and extra-renal phenotypes of Glis3 null mice prevented us from using these mice to study the potential functional interaction of Glis3 with polycystic kidney disease. Therefore, we generated a Glis3 conditional allele (*Glis3^fl^*) by using CRISPR/Cas9 to insert lox*P* sites flanking exon 6 to delete the fifth zinc finger domain (Supplementary Figure 1). This has been shown to be sufficient to abolish the DNA binding ability of Glis3 and therefore the transcriptional activity of Glis3 [^16^]. We crossed the *Glis3^fl^* allele with *Pkd1^fl/fl^; Pax8^rtTA^; TetO^Cre^* mice to achieve doxycycline inducible deletion of target genes selectively in kidney tubule epithelial cells. We have previously shown that Glis2 upregulation is a reliable in vitro indicator of CDCA in vitro^9^. We began by examining whether *Glis3* inactivation affects polycystin-dependent Glis2 expression, in cultured kidney primary kidney cell cultures from mice with *Pkd1^fl/fl^; Pax8^rtTA^; TetO^Cre^* and *Glis3^fl/fl^; Pkd1^fl/fl^; Pax8^rtTA^; TetO^Cre^* genotypes without or with doxycycline treatment in vitro. RT-qPCR confirmed *Glis3* and *Pkd1* knockout in the corresponding samples (Figure 1b). *Glis2* mRNA levels were significantly increased following inactivation of *Pkd1* (Figure 1b, *left*) and remained elevated in *Glis3* and *Pkd1* double mutants (Figure 1b, *right*). Glis2 protein expression was also unchanged in *Glis3* and *Pkd1* double mutant cells compared to *Pkd1* single knockouts (Figure 1c). *Glis3* is unlikely to function as a regulator of *Glis2* expression in the setting of *Pkd1* inactivation.

**Figure 1.**
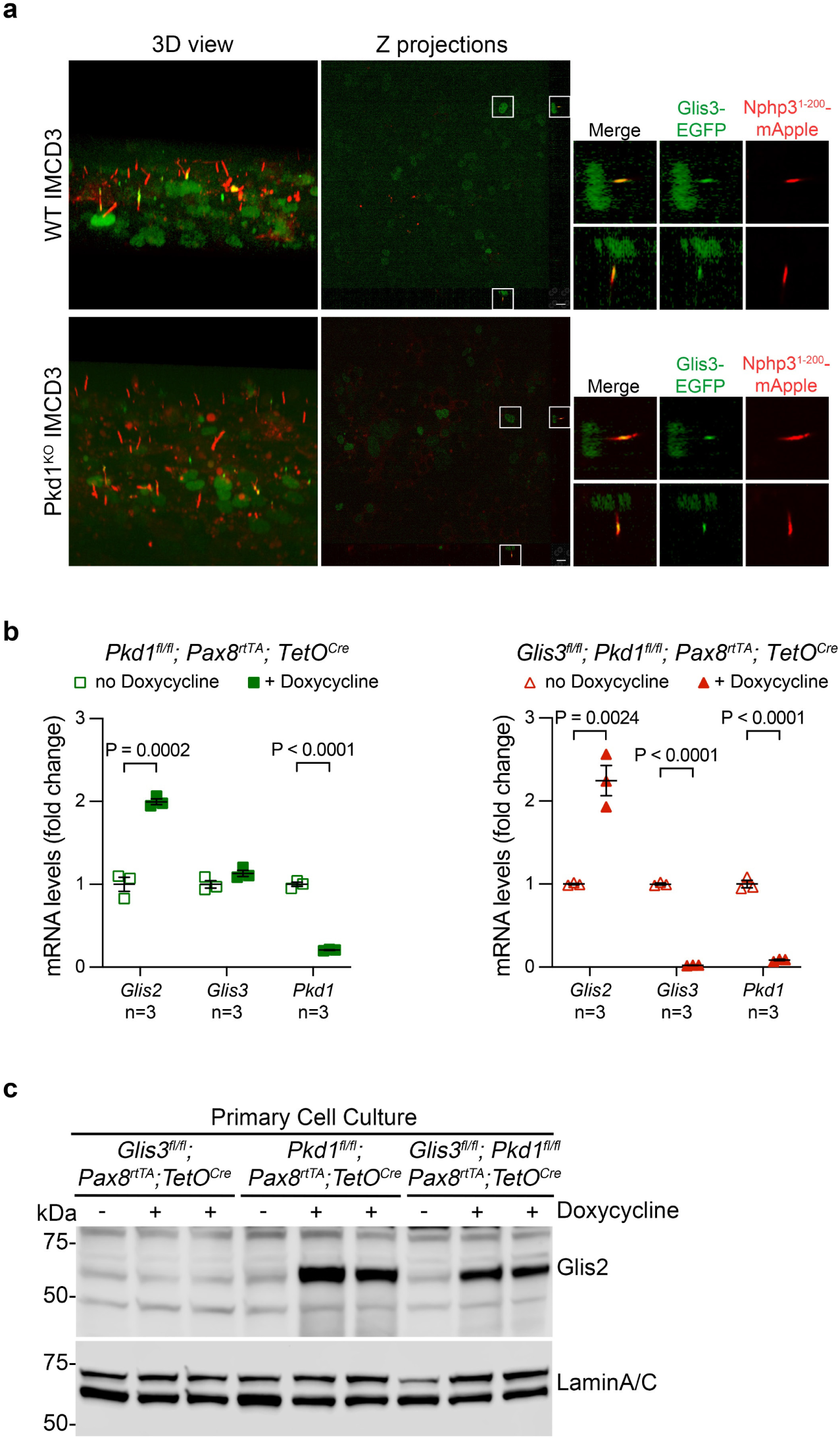
*Glis3* does not affect polycystin dependent *Glis2* expression. **a**, Isogenic WT or Pkd1^KO^ IMCD3 cells stably expressing Glis3-EGFP and the Nphp3^(1–200)^-mApple cilia marker cultured under conditions to form cilia and imaged live by confocal microscopy. 3D reconstructions and z-stack projections with magnified views of boxed regions. Scale bar, 10 μm. **b**, RT-qPCR of renal primary cell culture following in vitro doxycycline induced inactivation of either *Pkd1* alone or *Glis3* and *Pkd1* together. Fold change of each gene following doxycycline induction (closed symbol) is shown relative to the expression of same gene without doxycycline (open symbol), which is set to 1.0. Each data point is an independent primary cell culture from a single mouse with the indicated genotype. Statistical significance for each gene is determined by unpaired two-tailed Student’s t test and presented as mean ± s.e.m. **c**, Immunoblots of nuclear fraction of renal primary cell following in vitro doxycycline induced inactivation of either *Glis3* only, *Pkd1* only, or *Glis3* and *Pkd1*.

To determine the role of Glis3 in polycystic kidney disease in vivo, we generated mice with the following four genotypes: wildtype (WT), *Glis3^fl/fl^; Pax8^rtTA^; TetO^cre^* (Glis3^KO^), *Pkd1^fl/fl^; Pax8^rtTA^; TetO^cre^* (Pkd1^KO^), and *Glis3^fl/fl^; Pkd1^fl/fl^; Pax8^rtTA^; TetO^cre^*(Pkd1^KO^+Glis3^KO^). We started with the early-onset model where doxycycline in drinking water was administrated to the nursing dams from postnatal day 0 (P0) to postnatal day 14 (P14), and the experimental pups were examined at P14. Compared to the Pkd1^KO^ cystic controls, kidney-to-body weight ratio and cystic index were significantly increased in Pkd1^KO^+Glis3^KO^, indicating a significant worsening of the polycystic phenotype (Figure 2a-e, Supplementary Figure 2). Glis3^KO^ mice showed very occasional and sporadic tubule dilation (Figure 2b), but there was no significant abnormal phenotype in the P14 kidney (Figure 2c-e). All kidneys showed intact cilia detected by immunofluorescence microscopy using antibodies to acetylated α-tubulin (Figure 2f), confirming that Glis3 inactivation does not affect gross cilia structure. We next examined the role of Glis3 in adult-onset models of ADPKD where experimental animals were administrated doxycycline in drinking water from P28 to P42 and were examined at 14-weeks age for the polycystic phenotype. Similarly to the early-onset model, Pkd1^KO^+Glis3^KO^ mice at 14-weeks showed worsen polycystic kidney disease progression with significantly increased kidney-to-body weight ratio, cystic index, and blood urea nitrogen (BUN) level compared to Pkd1^KO^ (Figure 3a-e, Supplementary Figure 3). The 14 week endpoint was chosen because Pkd1^KO^+Glis3^KO^, but not Pkd1^KO^ mice have decreased survival after that timepoint. Of note, in distinction from the early-onset model, Glis3^KO^ kidneys at 14-weeks do not exhibit any histological changes (Figure 3b). Glis3^KO^ kidneys are also completely normal when they were further aged and evaluated at 18-weeks and 24-weeks (Figure 3f). These results indicate that the worsening phenotype observed in Pkd1^KO^+Glis3^KO^ is not due to additive effect of the two single mutants but suggests that Glis3 genetically interacts with *Pkd1*. *Glis3* is a modifier of cyst progression in models of ADPKD.

**Figure 2.**
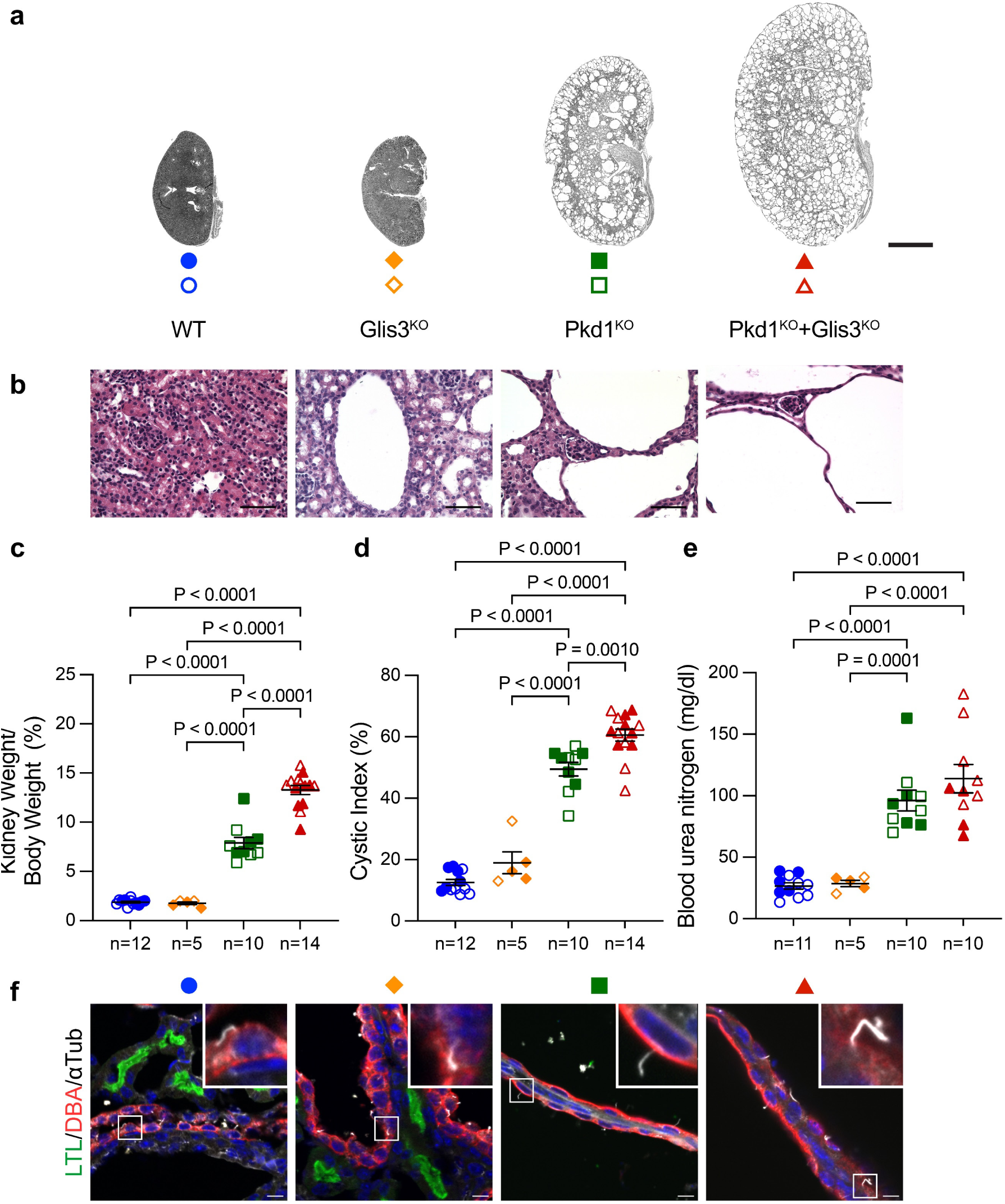
*Glis3* inactivation worsens ADPKD in an early model of *Pkd1*. **a**, Representative images of kidneys from mice at P14 with indicated genotypes. Oral doxycycline was administrated to the nursing dams from P0 to P14, and the kidneys of the pups were examined at P14. Scale bar, 2 mm. **b**, Representative images of hematoxylin and eosin (H&E) staining for the corresponding genotype. Scale bar, 50 μm. **c-e**, Aggregate quantitative data for kidney-to-body weight ratio (**c**), cystic index (**d**), and blood urea nitrogen (**e**). Colors and symbol shapes correspond to genotype defined in **a**. Sex of mice are shown for male (closed symbols) and female (open symbols). *n*, number of mice in each group. Multiple-group comparisons were performed by one-way ANOVA followed by Tukey’s multiple-comparison test, presented as mean ± s.e.m. **f**, Immunofluorescence of Lotus tetragonolobus lectin (LTL; proximal tubule), Dolichos biflorus agglutinin (DBA; collecting duct), and acetylated α-tubulin on cryosections from kidneys with genotypes indicated by color symbols in **a**. Boxed regions show magnified views. Scale bar, 10 μm.

**Figure 3.**
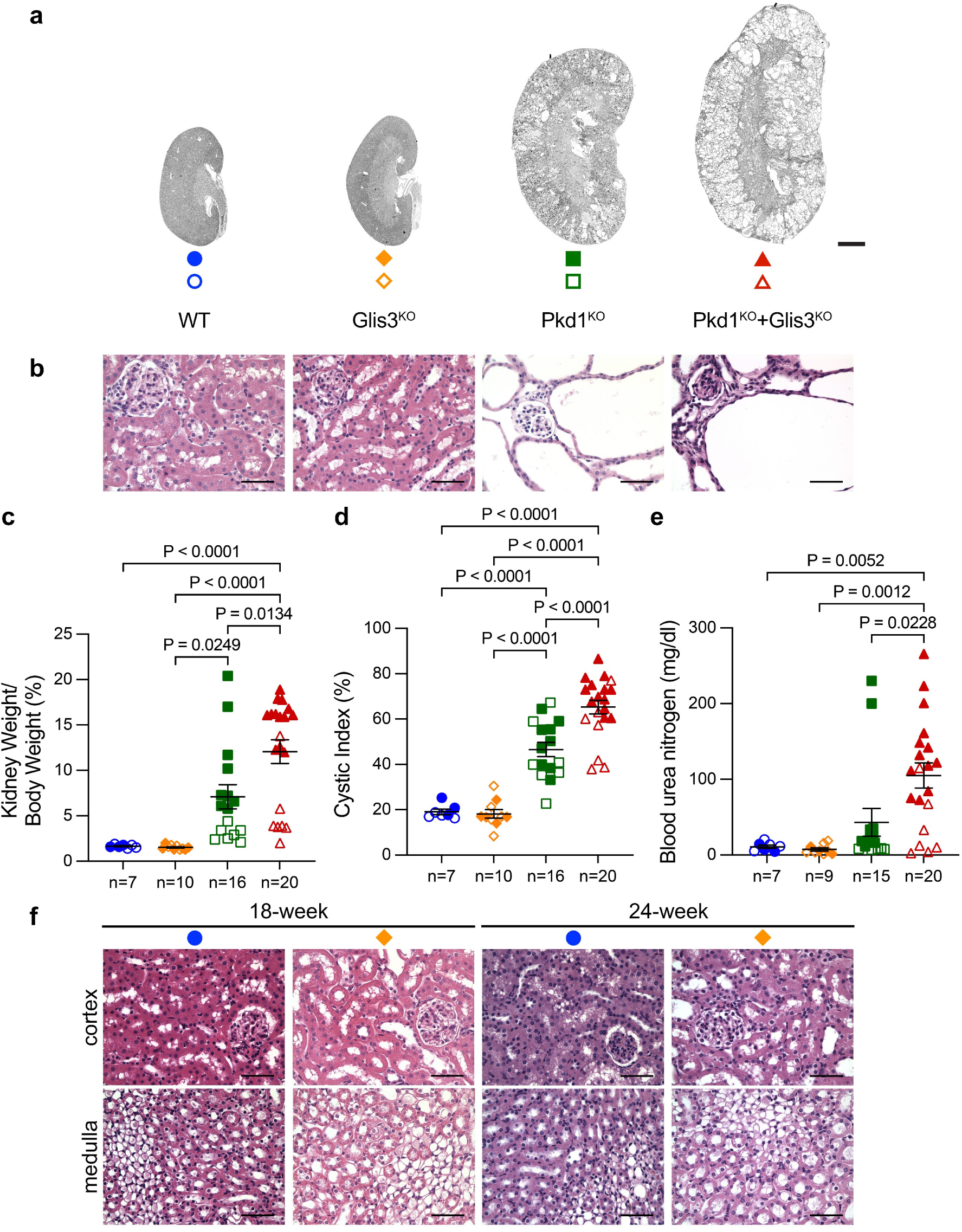
*Glis3* inactivation worsens ADPKD in an adult model of *Pkd1*. **a**, Representative images of kidneys from mice at 14-weeks of age with indicated genotypes. All mice were administrated with oral doxycycline from P28 to P42 and examined at 14-weeks. Scale bar, 2 mm. **b**, Representative images of H&E staining for the corresponding genotype. Scale bar, 50 μm. **c-e**, Aggregate quantitative data for kidney-to-body weight ratio (**c**), cystic index (**d**), and blood urea nitrogen (**e**). Colors and symbol shapes correspond to genotype defined in **a**. Sex is shown for male (closed symbols) and female (open symbols) mice. *n*, number of mice in each group. Multiple-group comparisons were performed by one-way ANOVA followed by Tukey’s multiple-comparison test, presented as mean ± s.e.m. **f**, Representative H&E staining images for WT and Glis3^KO^ mice aged to 18- and 24-weeks. Scale bar, 50 μm.

To begin to understand how Glis3 inactivation exacerbates cyst progression in ADPKD, we performed RNA-Seq and ATAC-Seq across the four genotypes: WT, Glis3^KO^, Pkd1^KO^, and Pkd1^KO^+Glis3^KO^. All samples were from male mice to limit confounding sex-dependent genomic effects and gene expression changes. All animals were induced with doxycycline from P28-P42. RNA and Tn5 transposase tagmented DNA was both prepared from the same kidneys at 7 weeks age (P49)—the same age used in our original TRAP RNA-Seq studies^9^. Principal component analysis (PCA) for the RNA-Seq showed promising clustering of samples by genotype (Figure 4a). Pkd1^KO^ and Pkd1^KO^+Glis3^KO^ clustered near each other, suggesting *Pkd1* inactivation is a major driver of the transcriptomic changes (Figure 4a). To determine whether the core transcriptomic signature of cyst initiation after *Pkd1* inactivation is affected by Glis3, we examined the differential expression of the previously published 73 CDCA pattern genes^9^ across the four genotypes. Heatmaps with unsupervised hierarchical clustering showed clear separation of genotypes that will not form cysts (WT and Glis3^KO^) from genotypes that will form cysts (Pkd1^KO^ and Pkd1^KO^+Glis3^KO^) (Figure 4b). These data confirm the specificity of the 73 gene CDCA translatome expression signature as an early in vivo bulk RNA-Seq readout for polycystic kidney disease before overt cyst formation. The CDCA gene pattern was not affected by Glis3^KO^ in Pkd1^KO^+Glis3^KO^, supporting the hypothesis that the effect of Glis3^KO^ on the polycystic kidney phenotype likely involves pathways other than CDCA (Figure 4b). We next examined differentially expressed genes (DEG) across the entire transcriptome by applying a false discovery rate of (FDR) ≤0.05 as the significance threshold. To examine the Glis3 dependent transcriptome changes relative to both WT and Pkd1^KO^ conditions, we compared Glis3^KO^ to WT and Pkd1^KO^+Glis3^KO^ to Pkd1^KO^. This yielded 2404 and 537 DEGs, respectively (Table 1). Gene ontology analyses of the 2404 DEG show strong signals for changes in metabolic processes associated with Glis3^KO^ alone (Figure 4c). The 537 DEG in the Pkd1^KO^+Glis3^KO^ to Pkd1^KO^ comparison, presumably enriched for the genes most responsible for the phenotypic worsening due to Glis3^KO^, highlighted a striking enrichment of Fatty Acid (FA) Metabolic Process (Figure 4d). FA metabolism process genes in Pkd1^KO^+Glis3^KO^ showed a significant skew toward downregulation relative to Pkd1^KO^ (42 downregulated, 18 upregulated; *P* <1.3e^-3^ by exact binomial test; Figure 4e). Gene set enrichment analysis in Pkd1^KO^+Glis3^KO^ compared to Pkd1^KO^ confirmed the strong enrichment for FA metabolism and predominance of downregulated genes (Figure 4f). Dysregulated FA metabolism is a compelling candidate process for contributing to the worsening of polycystic kidney disease in Pkd1^KO^+Glis3^KO^ mouse models compared to Pkd1^KO^ alone.

**Figure 4.**
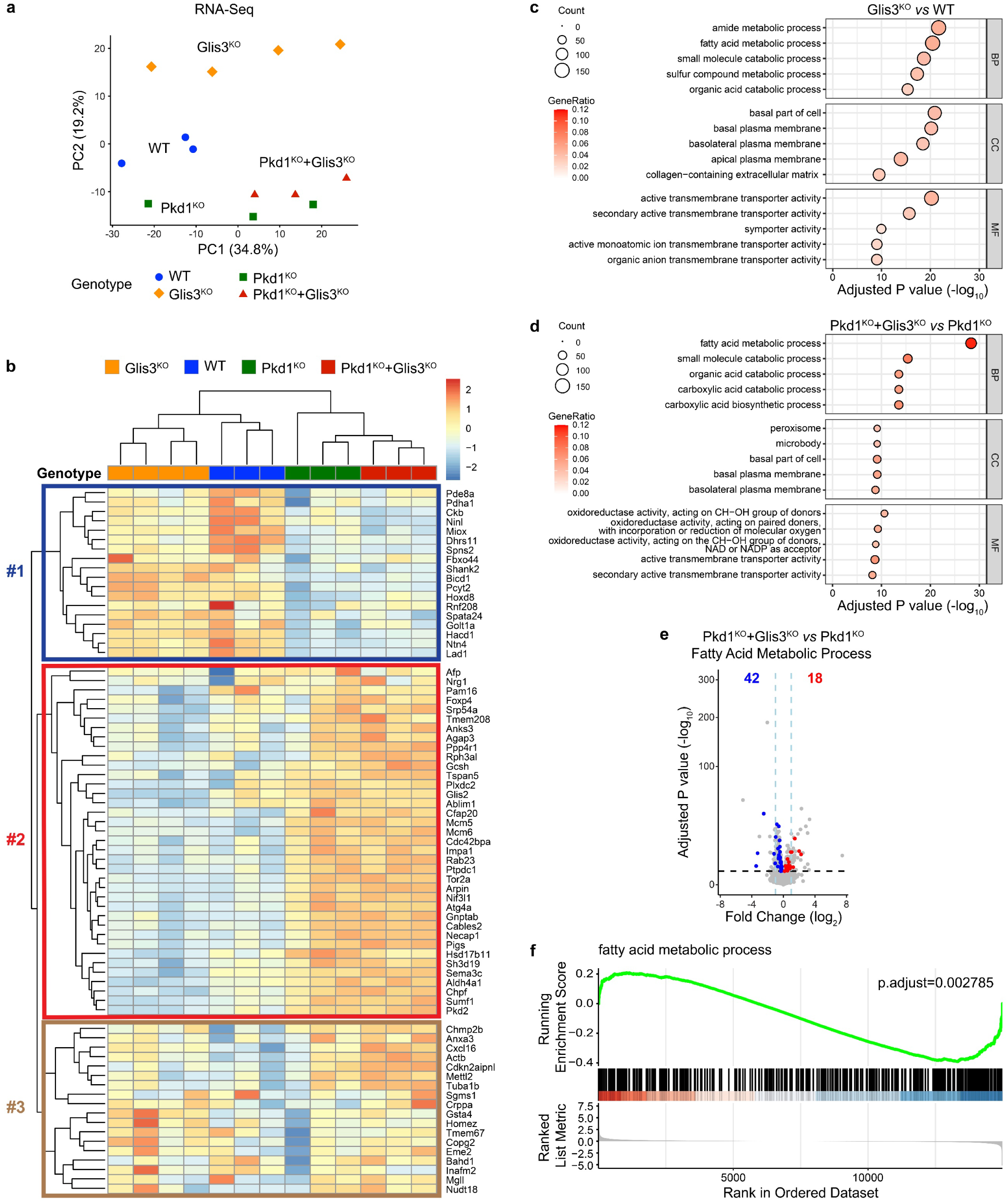
Dysregulated metabolism gene expression in *Glis3* inactivation. **a**, PCA plot of RNA-Seq showing the global transcriptomic difference among the indicated genotypes. Each symbol represents a biological replicate. **b**, Heatmap with unsupervised hierarchical clustering showing relative gene expression of the 73 CDCA signature genes with the indicated genotypes. **c,d**, Gene ontology analysis performed using the *clusterProfiler* R package with gene lists with FDR ≤0.05 threshold for significance in Glis3^KO^ compared to WT (**c**) and Pkd1^KO^+Glis3^KO^ compared to Pkd1^KO^ (**d**). BP, biological process; CC, cellular component; MF, molecular function. **e**, Volcano plot showing the expression of the 60 genes in the GO term “Fatty acid metabolic process” in Pkd1^KO^+Glis3^KO^ to Pkd1^KO^ comparison. Red, upregulated; blue, downregulated. **f**, Gene set enrichment analysis highlighting the strong enrichment for FA metabolism in Pkd1^KO^+Glis3^KO^ compared to Pkd1^KO^.

**Table 1.** Numbers of differentially expressed genes (DEG) and differentially accessible regions (DAR) in pairwise comparisons of RNA-Seq and ATAC-Seq.

| Pairwise comparison | RNA-Seq<br>DEGs | ATAC-Seq DARs | | Number of DARs<br>(FDR $\leq 0.05$ & Promoter)<br>overlapped with DEGs (FDR $\leq 0.05$) | |
| --- | --- | --- | --- | --- | --- |
| | FDR $\leq 0.05$ | FDR $\leq 0.05$ | FDR $\leq 0.05$ &<br>Promoter | Total | Same Direction<br>Change |
| Glis3<sup>KO</sup> vs WT | 2404 | 3776 | 697 | 234 | 216<br>( 26 / 190 ) |
| Pkd1<sup>KO</sup> vs WT | 1361 | 3147 | 445 | 68 | 62<br>( 23 / 39 ) |
| Pkd1<sup>KO</sup>+Glis3<sup>KO</sup> vs WT | 2053 | 3679 | 499 | 91 | 73<br>( 27 / 46 ) |
| Pkd1<sup>KO</sup>+Glis3<sup>KO</sup> vs Pkd1<sup>KO</sup> | 537 | 330 | 44 | 6 | 6<br>( 2 / 4 ) |

Table showing numbers of DEG and DARs identified in the indicated pairwise comparisons. In the rightmost columns, DARs in promoter regions are overlapped with DEGs and filtered for same direction change. Red, more accessible; blue, less accessible.

We next sought to integrate the RNA-Seq and ATAC-Seq data obtained from the same biological samples. PCA of ATAC-Seq showed clustering of samples by genotype (Figure 5a). We observed strong signal-to-noise at transcription start sites (TSS) across all samples (Figure 5b), highlighting successful enrichment of open chromatin accessibility at promoter-proximal regions. In total, we detected 129,765 consensus peaks, of which 21.49% were annotated to promoter regions defined as ±2 kb from TSS and 33.45% were mapped to distal intergenic regions (Figure 5c). This distribution aligns with a previous report of transcriptional regulatory element localization in the kidney^29^. We identified differentially accessible chromatin regions (DAR) in pairwise comparisons among the four genotypes with FDR ≤0.05 as the threshold for significance (Table 1). We further refined this to select DAR located in the promoter regions near TSS that also overlapped with DEG identified in the respective pairwise genotype group comparisons for RNA-Seq (Table 1). Within these overlap groups, we further constrained analysis to DAR in TSS regions that overlapped with DEG with the same direction of change, meaning that more accessible TSS region DAR in a pairwise comparison had upregulated genes and less accessible TSS region DAR had downregulated genes. For the Glis3^KO^ compared to WT, 234 out of 697 DAR in the region of TSS overlapped DEG from the corresponding RNA-Seq comparison. Of these, 216 DAR (>90%), had the same direction change as the corresponding 190 DEG, (Table 1, Figure 5d). There were fewer DEG than DAR because several genes has more than one TSS region DAR with same direction change associated with it. By comparison, Pkd1^KO^ compared to WT had 62 of 68 DAR overlapping with 55 DEG in have same direction change (Table 1, Figure 5e). Notably, these 55 DEG included *Lad1*, *Ntn4* and *Spns2*, three genes significantly downregulated by Pkd1 inactivation in the TRAP RNA-Seq^9^. All three genes showed downregulated gene expression and less accessible chromatin in the TSS region in Pkd1^KO^ compared to WT. Overall, these data suggest an informative correlation between overlapping changes chromatin accessibility in TSS regions and gene expression from bulk RNA-Seq.

**Figure 5.**
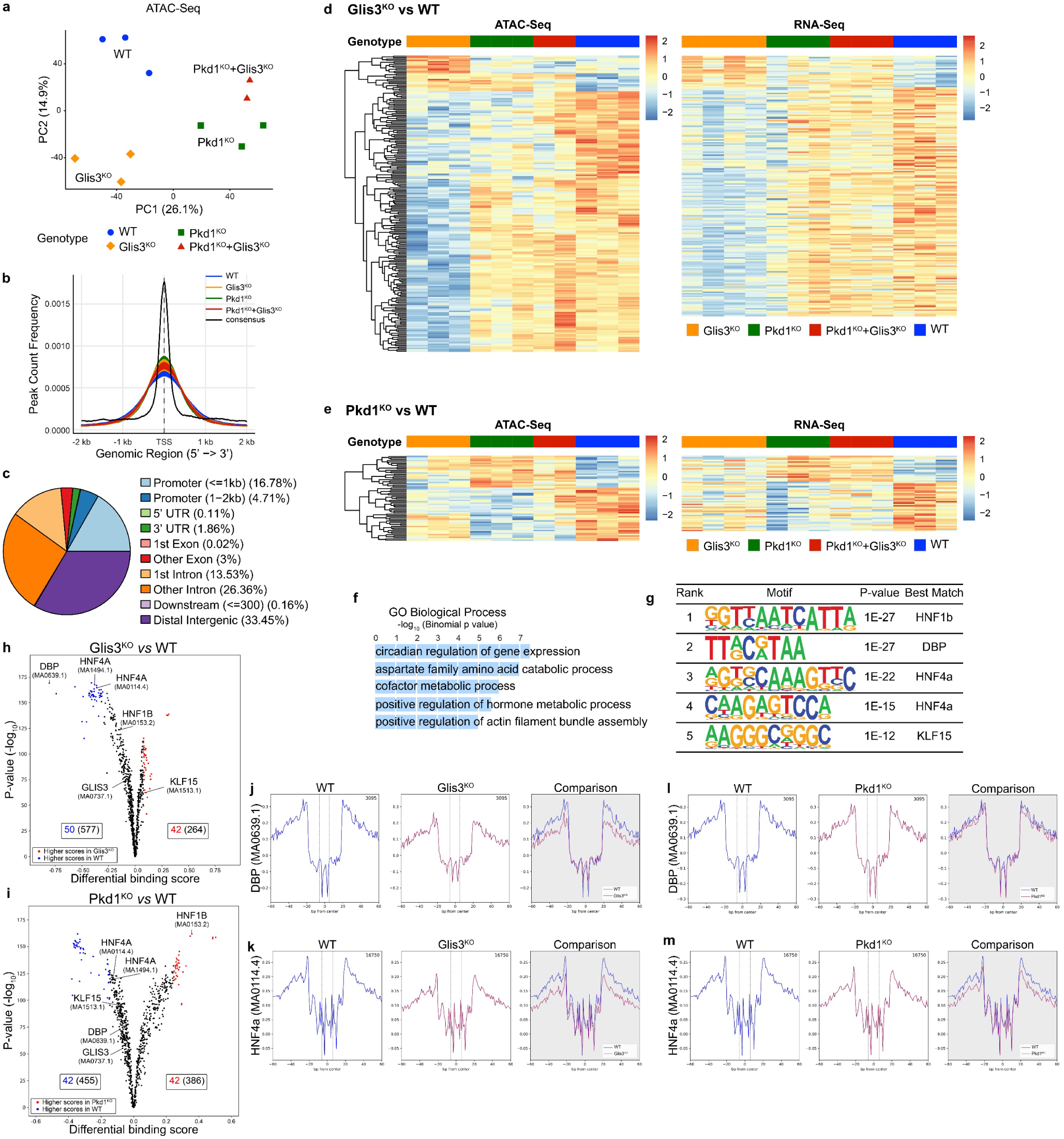
Genomic accessibility and transcription factor footprinting in Glis3 inactivation. **a**, PCA plot of ATAC-Seq showing the global chromatin accessibility differences among the indicated genotypes. Each symbol is a biological replicate. **b**, Transcription start site (TSS) enrichment profile of ATAC-Seq samples. The aggregate distributions of ATAC-Seq consensus peaks for each genotype are shown relative to TSS and flanking regions spanning ±2 kb upstream and downstream of TSS. **c**, Genomic annotation of consensus peaks. 21.49% of the peaks fall into promoter regions. **d**,**e**, Heatmaps show hierarchical clustering for fold change in DAR grouped by genotype (*left panels*); DEG from RNA-Seq (*right panels*) are shown in the order defined by their respective DAR hierarchical clustering. **d**, Heatmaps illustrating the correlation of 216 ATAC-Seq DAR peaks in promoter regions and overlapping 190 RNA-Seq DEG with the same direction change in expression in Glis3^KO^ *vs* WT. **e**, Heatmaps of 62 ATAC-Seq DAR peaks in promoter regions and overlapping 55 RNA-Seq DEG with the same direction change in Pkd1^KO^ *vs* WT. **f**, Bar plot showing gene ontology analysis of the 216 gene overlap with same direction change in Glis3^KO^ *vs* WT comparison using the Genomic Regions Enrichment of Annotations Tool (GREAT) tool. **g**, Significant result of *de novo* motif analysis for the 190 downregulated TSS region DAR overlapped with DEG in Glis3^KO^ *vs* WT. **h,i**, Volcano plots showing differential binding score distribution of the 841 transcription factor binding sites scanned by TOBIAS in Glis3^KO^ *vs* WT (**h**) and Pkd1^KO^ *vs* WT (**i**). Blue, less occupancy in Glis3^KO^ (**h**) or Pkd1^KO^ (**i**); red, more occupancy in Glis3^KO^ (**h**) or Pkd1^KO^ (**i**). **j-m**, Footprint comparisons for transcription factors Dbp (MA0639.1; **j,l**) and Hnf4a (MA0114.4; **k,m**) in Glis3^KO^ *vs* WT (**j,k**) and Pkd1^KO^ *vs* WT (**l,m**).

We next explored the potential functional significance of the 216 DAR in the Glis3^KO^ *vs* WT comparison that showed overlap with DEG to gain insight into the consequences of the genomic changes following Glis3 inactivation. These 216 DAR had a significant preference for less accessible chromatin (190 *vs* 26; *P* <2.2e^-16^ by exact binomial test; Table1), suggesting that Glis3 is an activating transcription factor that normally promotes chromatin accessibility. Analysis using the Genomic Regions Enrichment of Annotations Tool (GREAT)^34^ suggested that the cis functions of the 216 DAR are related to circadian regulation and metabolic processes (Figure 5f). De novo motif analysis with HOMER for the 190 DAR that were less accessible in Glis3^KO^ yielded significant signals for only four transcription factors: the two kidney epithelial related transcription factors, Hnf1b^37,38^ and Hnf4a^39–41^; the circadian related transcription factor, Dbp^42,43^; and the metabolism and circadian related transcription factor, Klf15 [^42,44,45^] (Figure 5g; Supplementary Figure 4). Glis3^KO^ results in downregulation of chromatin access for epithelial and circadian related transcriptional programs.

Footprinting analysis in ATAC-Seq provides information about transcription factor occupancy at binding motif sites across the genome. We applied TOBIAS (Transcription factor Occupancy prediction By Investigation of ATAC-seq Signal)^35^ to detect transcription factor footprints and infer transcription factor binding from the ATAC-Seq data. Of 841 transcription factor binding sites scanned by TOBIAS from the JASPAR database^46^, 577 trended toward reduced occupancy and 264 showed increased occupancy in Glis3^KO^ relative to WT (Figure 5h). Pkd1^KO^ showed more balanced trends with 455 toward reduced and 386 toward increased occupancy (Figure 5i). These data show that Glis3^KO^ not only results in less accessible chromatin (Table 1, Figure 5d), but also results in reduced transcription factor occupancy of motif binding sites (Figure 5h), reinforcing the hypothesis that Glis3 normally functions as an activating transcription factor. The top 5% footprint scores assigned by TOBIAS represent high confidence motif binding sites in the dataset^35^. Glis3^KO^ had 50 less occupied and 42 more occupied instances meeting this confidence threshold (Figure 5h). Among these, Dbp showed the greatest differential binding score (−0.8054) and the two binding motifs for Hnf4a also showed greatly reduced occupancy in Glis3^KO^ compared to WT (Figure 5h,j,k and Supplementary Figure 5a). Neither of the other two motifs identified by the de novo motif analysis, Hnf1b and Klf15 (Figure 5g), were present in the 95% quantile in the footprinting analysis (Figure 5h). By comparison, footprinting analysis in Pkd1^KO^ did not show significant binding motif occupancy changes for Dbp, Hnf4a or Klf15 compared to WT (Figure 5i,l,m and Supplementary Figure5b), but Hnf1b showed one of the most significant increases in binding motif occupancy in Pkd1^KO^ (Figure 5i). We also evaluated the footprinting for the three published Glis3 binding motifs^33,46,47^. The Glis3 motifs from either the JASPAR or HOMER databases did not show footprints of transcription factor occupancy in WT or Glis3^KO^ (Figure 5h and Supplementary Figure 5c,d). The footprint motif for Glis3 identified by a recent ChIP-Seq study^47^ was readily identified, but there was no difference in binding occupancy for this motif when comparing Glis3^KO^ to WT (Supplementary Figure 5e). Glis3^KO^ showed overall reduced transcription factor binding occupancy that included significant reductions in binding by Dbp and Hnf4a.

Finally, we integrated ATAC-Seq DAR data for Glis3^KO^ *vs* WT with the recent Glis3 ChIP-Seq dataset^47^. We identified 371 of 3776 DARs (9.80%) that overlapped with Glis3 ChIP genomic peaks. This suggests that Glis3 may in large part affect transcriptional regulation through coordination with other transcription factors, rather than changing chromatin accessibility by directly binding to DNA. To explore potential direct targets of Glis3, we intersected the ChIP-Seq peaks that fall into the promoter regions with the 190 downregulated DAR in promotor regions that overlapped with DEG in Glis3^KO^ vs. WT. We identified 23 genes (Supplementary Figure 6) which are transcriptionally downregulated and have less open chromatin in the promoter region in Glis3^KO^ *vs* WT, and whose promoter directly bind to Glis3 based on published ChIP-Seq data. This overlapping dataset gene list is notable for the circadian transcription factor Dbp. Dbp and perhaps other genes on this list are likely candidates for direct regulation by Glis3 transcription factor activity.

## Discussion

*Gis2* is a necessary effector of the cyst promoting CDCA pathway downstream of polycystin inactivation in the setting of intact cilia. Based on this we tested a hypothesis, analogous to the functioning of Patched and the Gli transcription factors in Hedgehog signaling, that the Glis3 transcription factor located in cilia functions as the polycystin and cilia dependent upstream modulator of *Glis2* upregulation in ADPKD. We found this not be the case. *Glis3* inactivation did not change *Glis2* expression or expression of other CDCA pattern genes^9^. Notably, adult inactivation of *Glis3* selectively in kidney tubules did not lead to cyst formation as has been reported for *Glis3* germline mutant mice^16,17,48^. *Glis3* inactivation did, however, interact genetically with *Pkd1* to exacerbate the polycystic kidney disease phenotype in ADPKD mouse models based on selective kidney tubule inactivation of both genes. We propose that this effect of Glis3 is related to activity in pathways other than CDCA.

We applied an integrated analysis of in vivo transcriptomic and chromatin accessibility studies to understand the effects of loss of *Glis3* in kidney tubules in the presence and absence of *Pkd1* inactivation. A key feature to interpreting the results from the current study is that *Glis3* or *Pkd1* inactivation was confined to kidney tubule cells of the mature kidney and both RNA-Seq and ATAC-Seq were done from the same biological samples at a stage when there was effective tubule-cell autonomous inactivation of *Glis3* or *Pkd1* but no overt secondary phenotypes in the kidney or systemically^9^. In this setting, Glis3 inactivation largely resulted in decreased chromatin accessibility in promoter regions and this was associated with downregulation of gene expression from those regions. While the activating role of wild type Glis3 expression implied by this knockout result is consistent with ChIP-Seq studies^47^, the reduction in chromatin accessibility largely occurred in regions where Glis3 was not known to bind. Glis3 may therefore regulate promoter regions indirectly by interaction with other transcription factors or by association with enhancer or other genomic sites remote from promoter regions. The transcriptomic effects of *Glis3* inactivation in our RNA-Seq from normal adult kidneys showed strong association with metabolic processes and these effects appear to be related to differential expression of transmembrane transporter activities. This is generally consistent with a recent study of *Glis3* germline knockout mice that showed suppression of genes critical for mitochondrial biogenesis, OXPHOS, fatty acid oxidation, and the tricarboxylic acid (TCA) cycle^47^. Dual inactivation of *Glis3* with *Pkd1* accentuated a strong signal for downregulation of fatty acid metabolism processes. *Pkd1* mutant kidneys have defects in fatty acid oxidation and exacerbations of these defects likely worsen the progression of ADPKD^49^. Dysregulation of fatty acid metabolism by *Glis3* inactivation is the strongest candidate biologic process from our study to explain the worsening of the *Pkd1* ADPKD phenotype with *Glis3* inactivation.

*Hnf1b* and *Hnf4a* encode important transcription factors for kidney epithelial differentiation and function^40,50^. Motifs for both were identified in our ATAC-Seq and showed reduced chromatin accessibility in Glis3^KO^; however, Glis3 inactivation was most significantly associated with reduction of Hnf4a binding occupancy. The data suggest a context dependent role in the kidney for Glis3 as part of a transcriptional network which affects the accessibility of sites typically occupied by the Hnf1b and Hnf4a. Downregulation of *Hnf4a* has been shown to be a disease modifier associated with more rapid progression of ADPKD in vivo^51^. *HNF4a* knockout in human kidney organoids is associated with downregulation of lipid metabolism^39^ and dual inactivation of *Hnf4a* and *Pkd1* exacerbates polycystic kidneys in mice compared to Pkd1^KO^ alone^51^. Our study identifies Hnf4a accessible regions as a significant target of *Glis3* inactivation that was not previously known^47^. This Hnf4a related function may be the mechanistic connection with the alterations in lipid metabolism associated with worsening ADPKD in *Glis3* and *Pkd1* dual inactivation kidneys. The *de novo* transcription factor binding sites analysis for Glis3 also identified Dbp and Klf15. Of these, the signal for Dbp was the most significant across our analyses and overlapped with direct Glis3 binding data identified from ChIP-Seq^47^. Dbp is a transcription factor whose expression is controlled by the circadian clock genes and whose function is to effect transcriptional responses to circadian rhythm^42,52–54^. Recent data suggests disruption of circadian clocks may accelerate progression in ADPKD^55^, so the effect of *Glis3* on ADPKD progression may also be mediated by the downregulation of *Dbp*. *Glis3* may indirectly affect kidney metabolic programs including fatty acid oxidation through Hnf4a and kidney circadian programs through Dbp. The consequent metabolic and circadian perturbations are how loss of *Glis3* results in worsening of ADPKD severity.

## Acknowledgements

This work was supported by NIH/National Institute of Diabetes and Digestive and Kidney Diseases grants (nos. R01 DK120911, R01 DK100592, and RC2 DK120534 to S.S.) and a grant from the Amy P. Goldman Foundation to S.S. We are grateful for the generous support from Mr. and Mrs. Robert Roth.

## Author Contributions

Z.W. designed and performed experiments, analyzed data, and drafted figures and manuscript. This work was performed as part of the fulfilment of doctoral thesis requirements for Z.W.

J.G., supervised by H.Z., performed bioinformatic analyses.

X.T. and C.Z. provided reagents.

S.S. conceived the study, designed experiments, supervised the study, and wrote the manuscript.

## Data Sharing Statement

All the raw sequencing data and processed data have been deposited in the Gene Expression Omnibus (GEO) with the following accession number: GSE299238.

## Literatrue Cited

1. The European Polycystic Kidney Disease C. The polycystic kidney disease 1 gene encodes a 14 kb transcript and lies within a duplicated region on chromosome 16. Cell. 1994/06/17/ 1994;77(6):881–894. doi:10.1016/0092-8674(94)90137-6

2. TIPKD C. Polycystic kidney disease - the complete structure of the PKD1 gene and its protein. Cell. Apr 1995;81(2):289–298.

3. Mochizuki T, Wu G, Hayashi T, et al. PKD2, a gene for polycystic kidney disease that encodes an integral membrane protein. Article. Science. 1996;272(5266):1339–1342.

4. Bergmann C, Guay-Woodford LM, Harris PC, Horie S, Peters DJM, Torres VE. Polycystic kidney disease. Nat Rev Dis Primers. Dec 6 2018;4(1):50. doi:10.1038/s41572-018-0047-y

5. Harris PC, Torres VE. Genetic mechanisms and signaling pathways in autosomal dominant polycystic kidney disease. Review. J Clin Invest. Jun 2014;124(6):2315–2324. doi:10.1172/jci72272

6. Qiu JH, Germino GG, Menezes LF. Mechanisms of Cyst Development in Polycystic Kidney Disease. Adv Kidney Dis Heal. May 2023;30(3):209–219. doi:10.1053/j.akdh.2023.03.001

7. Padovano V, Podrini C, Boletta A, Caplan MJ. Metabolism and mitochondria in polycystic kidney disease research and therapy. Nat Rev Nephrol. Nov 2018;14(11):678–687. doi:10.1038/s41581-018-0051-1

8. Ma M, Tian X, Igarashi P, Pazour GJ, Somlo S. Loss of cilia suppresses cyst growth in genetic models of autosomal dominant polycystic kidney disease. Article. Nature Genet. Sep 2013;45(9):1004-+. doi:10.1038/ng.2715

9. Zhang C, Rehman M, Tian X, et al. Glis2 is an early effector of polycystin signaling and a target for therapy in polycystic kidney disease. Nat Commun. May 1 2024;15(1):3698. doi:10.1038/s41467-024-48025-6

10. Zhang F, Jetten AM. Genomic structure of the gene encoding the human GLI-related, Kruppel-like zinc finger protein GLIS2. Gene. Dec 12 2001;280(1-2):49–57.

11. Kim YS, Lewandoski M, Perantoni AO, Kurebayashi S, Nakanishi G, Jetten AM. Identification of Glis1, a novel Gli-related, Kruppel-like zinc finger protein containing transactivation and repressor functions. J Biol Chem. Aug 2002;277(34):30901–30913. doi:10.1074/jbc.M203563200

12. Kim YS, Nakanishi G, Lewandoski M, Jetten AM. GLIS3, a novel member of the GLIS subfamily of Kruppel-like zinc finger proteins with repressor and activation functions. Article. Nucleic Acids Res. Oct 2003;31(19):5513–5525. doi:10.1093/nar/gkg776

13. Chen LH, Chou CL, Knepper MA. A Comprehensive Map of mRNAs and Their Isoforms across All 14 Renal Tubule Segments of Mouse. Article. J Am Soc Nephrol. Apr 2021;32(4):897–912. doi:10.1681/asn.2020101406

14. Senee V, Chelala C, Duchatelet S, et al. Mutations in GLIS3 are responsible for a rare syndrome with neonatal diabetes mellitus and congenital hypothyroidism. Article. Nature Genet. Jun 2006;38(6):682–687. doi:10.1038/ng1802

15. London S, De Franco E, Elias-Assad G, et al. Case Report: Neonatal Diabetes Mellitus Caused by a Novel GLIS3 Mutation in Twins. Front Endocrinol. May 2021;12:8. 673755. doi:10.3389/fendo.2021.673755

16. Kang HS, Beak JY, Kim YS, Herbert R, Jetten AM. Glis3 Is Associated with Primary Cilia and Wwtr1/TAZ and Implicated in Polycystic Kidney Disease. Article. Mol Cell Biol. May 2009;29(10):2556–2569. doi:10.1128/mcb.01620-08

17. Watanabe N, Hiramatsu K, Miyamoto R, et al. A murine model of neonatal diabetes mellitus in Glis3-deficient mice. FEBS Lett. Jun 2009;583(12):2108–2113. doi:10.1016/j.febslet.2009.05.039

18. Beak JY, Kang HS, Kim YS, Jetten AM. Functional analysis of the zinc finger and activation domains of Glis3 and mutant Glis3(NDH1). Article. Nucleic Acids Res. Mar 2008;36(5):1690–1702. doi:10.1093/nar/gkn009

19. Shibazaki S, Yu Z, Nishio S, et al. Cyst formation and activation of the extracellular regulated kinase pathway after kidney specific inactivation of Pkd1. Hum Mol Genet. Jun 1 2008;17(11):1505–1516. doi:10.1093/hmg/ddn039

20. Shibazaki S, Yu ZH, Nishio S, et al. Cyst formation and activation of the extracellular regulated kinase pathway after kidney specific inactivation of Pkd1. Article. Hum Mol Genet. Jun 2008;17(11):1505–1516. doi:10.1093/hmg/ddn039

21. Chen S, Zhou Y, Chen Y, Gu J. fastp: an ultra-fast all-in-one FASTQ preprocessor. Bioinformatics. Sep 1 2018;34(17):i884–i890. doi:10.1093/bioinformatics/bty560

22. Dobin A, Davis CA, Schlesinger F, et al. STAR: ultrafast universal RNA-seq aligner. Bioinformatics. Jan 1 2013;29(1):15–21. doi:10.1093/bioinformatics/bts635

23. Liao Y, Smyth GK, Shi W. featureCounts: an efficient general purpose program for assigning sequence reads to genomic features. Bioinformatics. Apr 1 2014;30(7):923–930. doi:10.1093/bioinformatics/btt656

24. Love MI, Huber W, Anders S. Moderated estimation of fold change and dispersion for RNA-seq data with DESeq2. Genome Biol. 2014;15(12):550. doi:10.1186/s13059-014-0550-8

25. Yu G, Wang LG, Han Y, He QY. clusterProfiler: an R package for comparing biological themes among gene clusters. Omics. May 2012;16(5):284–287. doi:10.1089/omi.2011.0118

26. Buenrostro JD, Giresi PG, Zaba LC, Chang HY, Greenleaf WJ. Transposition of native chromatin for fast and sensitive epigenomic profiling of open chromatin, DNA-binding proteins and nucleosome position. Nat Methods. Dec 2013;10(12):1213-+. doi:10.1038/nmeth.2688

27. Buenrostro JD, Wu B, Chang HY, Greenleaf WJ. ATAC-seq: A Method for Assaying Chromatin Accessibility Genome-Wide. Current Protocols in Molecular Biology. 2015;109(1):21.29.21–21.29.29. 10.1002/0471142727.mb2129s109

28. Corces MR, Trevino AE, Hamilton EG, et al. An improved ATAC-seq protocol reduces background and enables interrogation of frozen tissues. Nat Methods. Oct 2017;14(10):959

29. Chen L, Chou CL, Yang CR, Knepper MA. Multiomics Analyses Reveal Sex Differences in Mouse Renal Proximal Subsegments. J Am Soc Nephrol. May 1 2023;34(5):829–845.

30. Langmead B, Salzberg SL. Fast gapped-read alignment with Bowtie 2. Nat Methods. Mar 4 2012;9(4):357–359. doi:10.1038/nmeth.1923

31. Zhang Y, Liu T, Meyer CA, et al. Model-based analysis of ChIP-Seq (MACS). Genome Biol. 2008;9(9):R137. doi:10.1186/gb-2008-9-9-r137

32. Yu G, Wang LG, He QY. ChIPseeker: an R/Bioconductor package for ChIP peak annotation, comparison and visualization. Bioinformatics. Jul 15 2015;31(14):2382–2383.

33. Heinz S, Benner C, Spann N, et al. Simple combinations of lineage-determining transcription factors prime cis-regulatory elements required for macrophage and B cell identities. Mol Cell. May 28 2010;38(4):576–589. doi:10.1016/j.molcel.2010.05.004

34. McLean CY, Bristor D, Hiller M, et al. GREAT improves functional interpretation of cis-regulatory regions. Nat Biotechnol. May 2010;28(5):495–501. doi:10.1038/nbt.1630

35. Bentsen M, Goymann P, Schultheis H, et al. ATAC-seq footprinting unravels kinetics of transcription factor binding during zygotic genome activation. Nat Commun. Aug 26 2020;11(1):4267. doi:10.1038/s41467-020-18035-1

36. Decuypere JP, Van Giel D, Janssens P, et al. Interdependent Regulation of Polycystin Expression Influences Starvation-Induced Autophagy and Cell Death. Int J Mol Sci. Dec 16 2021;22(24)doi:10.3390/ijms222413511

37. Ferrè S, Igarashi P. New insights into the role of HNF-1β in kidney (patho)physiology. Review. Pediatr Nephrol. Aug 2019;34(8):1325–1335. doi:10.1007/s00467-018-3990-7

38. Sánchez-Cazorla E, Carrera N, García-González MA. HNF1B Transcription Factor: Key Regulator in Renal Physiology and Pathogenesis. Review. Int J Mol Sci. Oct 2024;25(19):15. 10609. doi:10.3390/ijms251910609

39. Yoshimura Y, Muto Y, Omachi K, Miner JH, Humphreys BD. Elucidating the Proximal Tubule HNF4A Gene Regulatory Network in Human Kidney Organoids. J Am Soc Nephrol. Oct 1 2023;34(10):1672–1686. doi:10.1681/asn.0000000000000197

40. Marable SS, Chung E, Park JS. Hnf4a Is Required for the Development of Cdh6-Expressing Progenitors into Proximal Tubules in the Mouse Kidney. Journal of the American Society of Nephrology. Nov 2020;31(11):2543–2558. doi:10.1681/asn.2020020184

41. Martovetsky G, Tee JB, Nigam SK. Hepatocyte Nuclear Factors 4α and 1α Regulate Kidney Developmental Expression of Drug-Metabolizing Enzymes and Drug Transporters. Molecular Pharmacology. Dec 2013;84(6):808–823. doi:10.1124/mol.113.088229

42. Bingham MA, Neijman K, Yang CR, et al. Circadian gene expression in mouse renal proximal tubule. Am J Physiol Renal Physiol. Mar 1 2023;324(3):F301–f314. doi:10.1152/ajprenal.00231.2022

43. Costello HM, Johnston JG, Juffre A, Crislip GR, Gumz ML. Circadian clocks of the kidney : function, mechanism, and regulation. Physiological Reviews. Oct 2022;102(4):1669–1701. doi:10.1152/physrev.00045.2021

44. Piret SE, Attallah AA, Gu X, et al. Loss of proximal tubular transcription factor Krüppel-like factor 15 exacerbates kidney injury through loss of fatty acid oxidation. Kidney Int. Dec 2021;100(6):1250–1267. doi:10.1016/j.kint.2021.08.031

45. Rane MJ, Zhao Y, Cai L. Krϋppel-like factors (KLFs) in renal physiology and disease. EBioMedicine. Feb 2019;40:743–750. doi:10.1016/j.ebiom.2019.01.021

46. Rauluseviciute I, Riudavets-Puig R, Blanc-Mathieu R, et al. JASPAR 2024: 20th anniversary of the open-access database of transcription factor binding profiles. Nucleic Acids Res. Jan 5 2024;52(D1):D174–d182. doi:10.1093/nar/gkad1059

47. Collier JB, Kang HS, Roh YG, et al. GLIS3: A novel transcriptional regulator of mitochondrial functions and metabolic reprogramming in postnatal kidney and polycystic kidney disease. Mol Metab. Dec 2024;90:102052. doi:10.1016/j.molmet.2024.102052

48. Kang HS, Kumar D, Liao G, et al. GLIS3 is indispensable for TSH/TSHR-dependent thyroid hormone biosynthesis and follicular cell proliferation. J Clin Invest. Dec 1 2017;127(12):4326–4337. doi:10.1172/JCI94417

49. Menezes LF, Lin CC, Zhou F, Germino GG. Fatty Acid Oxidation is Impaired in An Orthologous Mouse Model of Autosomal Dominant Polycystic Kidney Disease. EBioMedicine. Mar 2016;5:183–192. doi:10.1016/j.ebiom.2016.01.027

50. Chan SC, Zhang Y, Shao A, et al. Mechanism of Fibrosis in HNF1B-Related Autosomal Dominant Tubulointerstitial Kidney Disease. J Am Soc Nephrol. 2018;29(10):2493–2509. doi:10.1681/asn.2018040437

51. Menezes LF, Zhou F, Patterson AD, et al. Network analysis of a Pkd1-mouse model of autosomal dominant polycystic kidney disease identifies HNF4α as a disease modifier. PLoS Genet. 2012;8(11):e1003053. doi:10.1371/journal.pgen.1003053

52. Zuber AM, Centeno G, Pradervand S, et al. Molecular clock is involved in predictive circadian adjustment of renal function. Proc Natl Acad Sci U S A. Sep 22 2009;106(38):16523–16528. doi:10.1073/pnas.0904890106

53. Jamadar A, Ward CJ, Remadevi V, et al. Circadian Clock Disruption and Growth of Kidney Cysts in Autosomal Dominant Polycystic Kidney Disease. J Am Soc Nephrol. 2024:10.1681/ASN.0000000528. doi:10.1681/asn.0000000528

54. Firsov D, Bonny O. Circadian rhythms and the kidney. Nature Reviews Nephrology. 2018/10/01 2018;14(10):626–635. doi:10.1038/s41581-018-0048-9

55. Jamadar A, Ward CJ, Remadevi V, et al. Circadian Clock Disruption and Growth of Kidney Cysts in Autosomal Dominant Polycystic Kidney Disease. J Am Soc Nephrol. Oct 14 2024;doi:10.1681/asn.0000000528

