## Supplementary Information for "Glis3 is a modifier of cyst progression in autosomal dominant polycystic kidney disease"

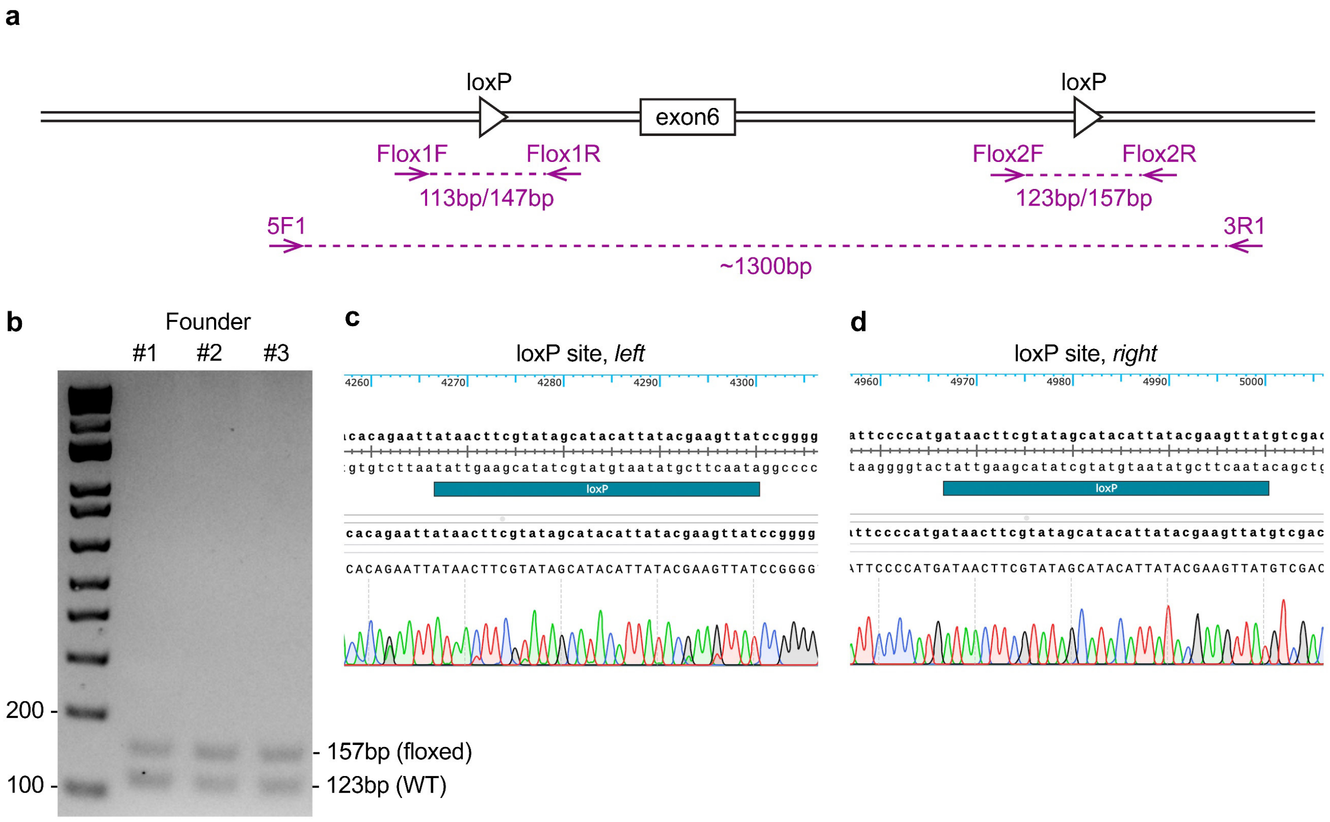


**Supplementary Figure 1. Generation of conditional *Glis3$^{fl}$* allele. a**, Schematic of *Glis3* conditional allele (*Glis3$^{fl}$*) and the location of genotyping primers. **b**, Representative agarose gel image with the Flox2F and Flox2R primer pair. **c,d**, Sequencing traces of the 5' (**c**) and 3' (**d**) loxP sites from founder mice #3.

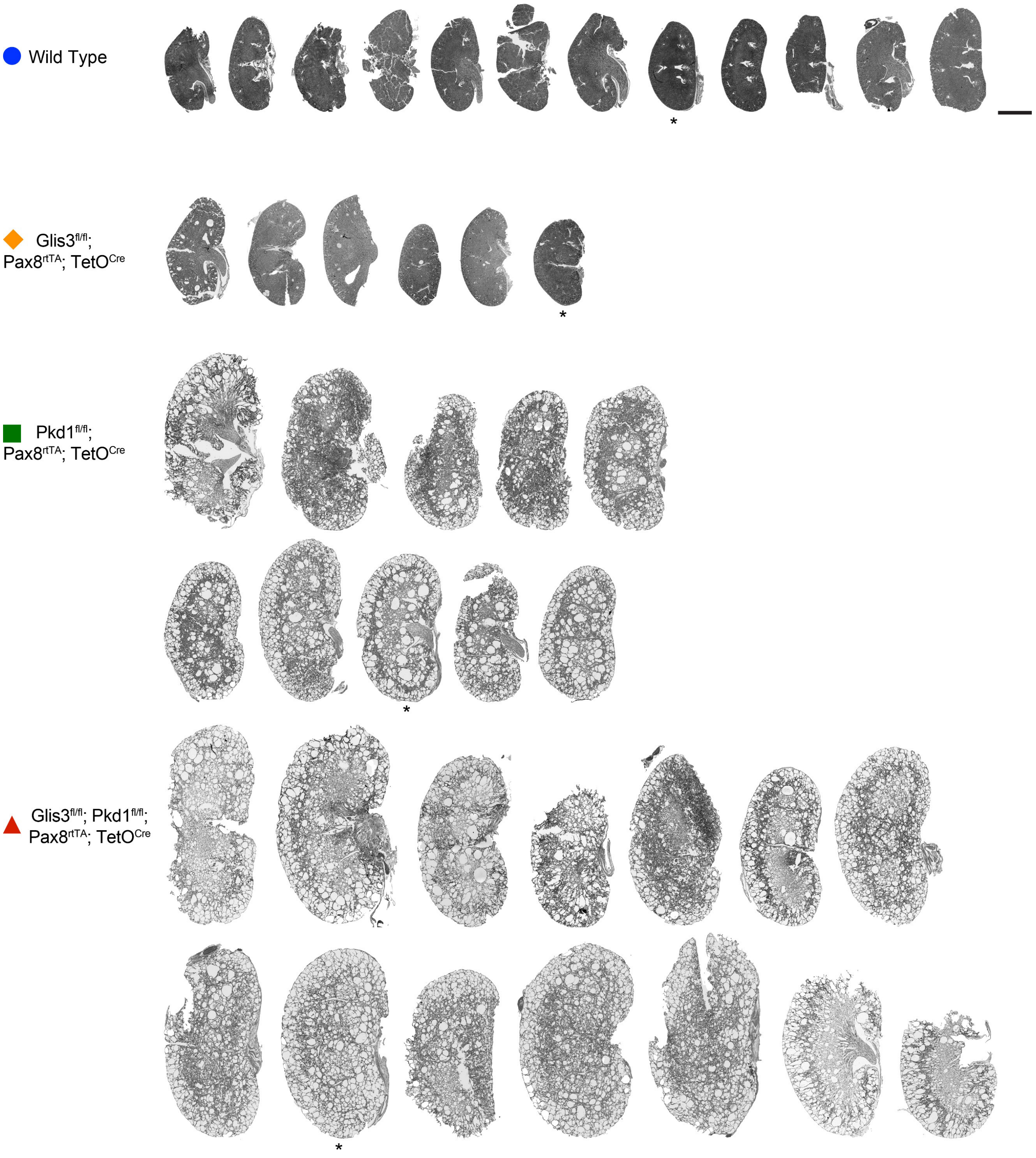


**Supplementary Figure 2. Images of all the histological sections used in Figure 2a-f.** Scale bar, 2 mm. Asterisks indicate the representative images used in Figure 2a.

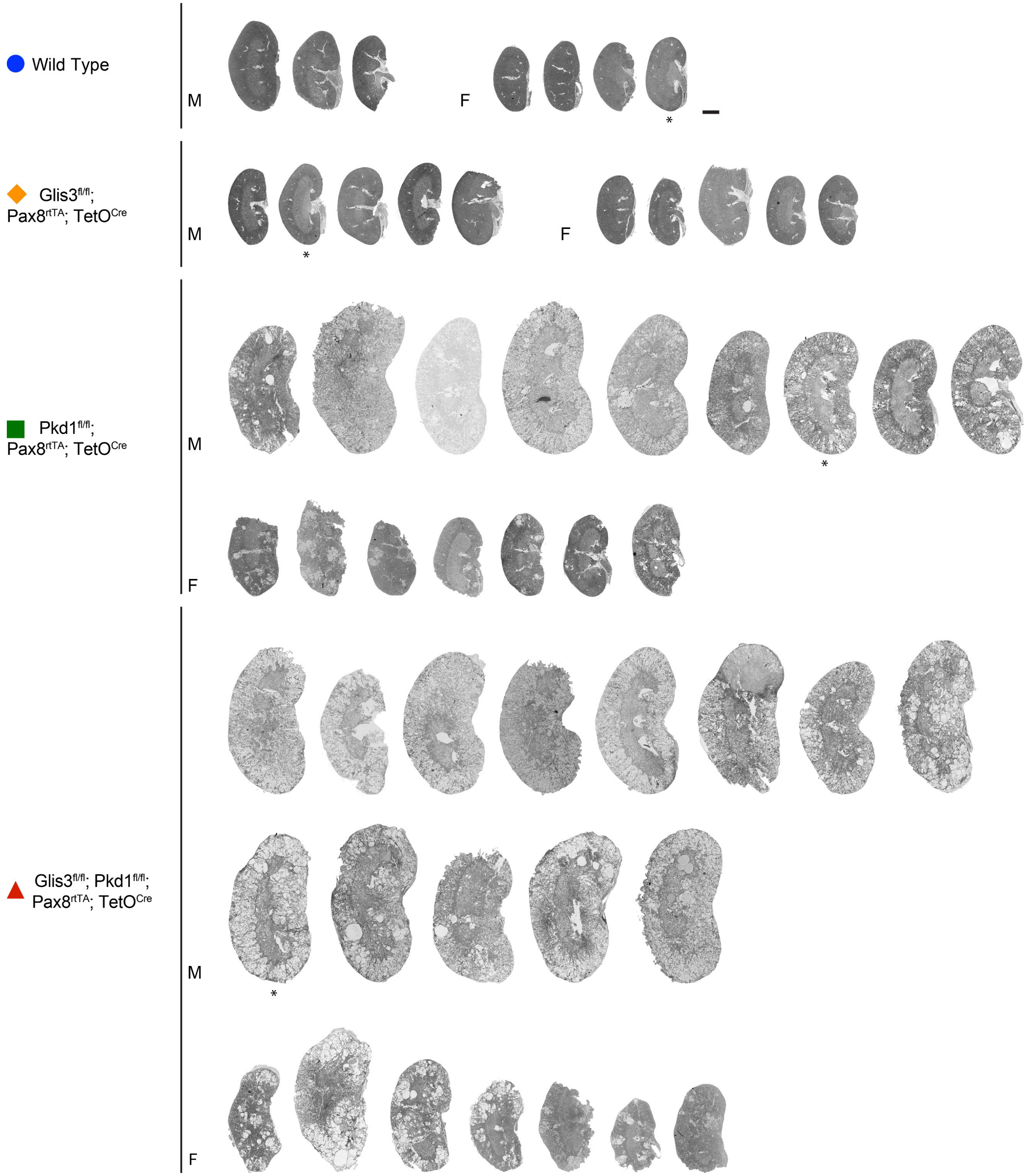


**Supplementary Figure 3. Images of all the histological sections used in Figure 3a-e.** Scale bar, 2mm. Asterisks indicate the representative images used in Figure 3a.

### Homer *de novo* Motif Results

Known Motif Enrichment Results
Gene Ontology Enrichment Results
If Homer is having trouble matching a motif to a known motif, try copy/pasting the matrix file into STAMP
More information on motif finding results: HOMER | Description of Results | Tips
Total target sequences = 179
Total background sequences = 48167
* - possible false positive

| Rank | Motif | P-value | log P-pvalue | % of Targets | % of Background | STD(Bg STD) | Best Match/Details | Motif File |
|---|---|---|---|---|---|---|---|---|
| 1 | 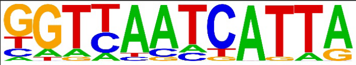 | 1e-27 | -6.443e+01 | 11.17% | 0.20% | 42.5bp (60.7bp) | HNF1b(Homeobox)/PDAC-HNF1B-ChIP-Seq(GSE64557)/Homer(0.887) More Information \| Similar Motifs Found | motif file (matrix) |
| 2 | 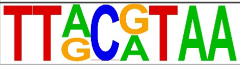 | 1e-27 | -6.408e+01 | 14.53% | 0.56% | 49.1bp (62.8bp) | DBP/MA0639.1/Jaspar(0.929) More Information \| Similar Motifs Found | motif file (matrix) |
| 3 | 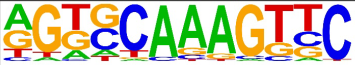 | 1e-22 | -5.243e+01 | 44.13% | 13.51% | 47.3bp (62.3bp) | HNF4a(NR),DR1/HepG2-HNF4a-ChIP-Seq(GSE25021)/Homer(0.913) More Information \| Similar Motifs Found | motif file (matrix) |
| 4 | 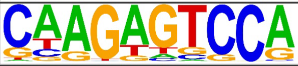 | 1e-15 | -3.494e+01 | 24.02% | 5.70% | 52.6bp (64.2bp) | PB0134.1_Hnf4a_2/Jaspar(0.777) More Information \| Similar Motifs Found | motif file (matrix) |
| 5 | 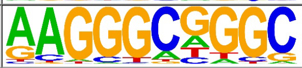 | 1e-12 | -2.927e+01 | 21.79% | 5.56% | 51.5bp (62.7bp) | KLF15/MA1513.1/Jaspar(0.769) More Information \| Similar Motifs Found | motif file (matrix) |
| 6 * | 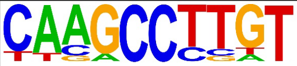 | 1e-10 | -2.395e+01 | 5.59% | 0.25% | 41.3bp (62.6bp) | ZFX(Zf)/mES-Zfx-ChIP-Seq(GSE11431)/Homer(0.645) More Information \| Similar Motifs Found | motif file (matrix) |
| 7 * | 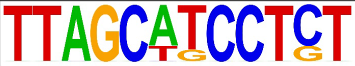 | 1e-9 | -2.141e+01 | 2.79% | 0.02% | 54.0bp (56.6bp) | Ets1-distal(ETS)/CD4+-PolII-ChIP-Seq(Barski_et_al.)/Homer(0.627) More Information \| Similar Motifs Found | motif file (matrix) |
| 8 * | 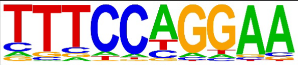 | 1e-9 | -2.138e+01 | 29.05% | 11.86% | 51.6bp (63.7bp) | Bcl6(Zf)/Liver-Bcl6-ChIP-Seq(GSE31578)/Homer(0.954) More Information \| Similar Motifs Found | motif file (matrix) |
| 9 * | 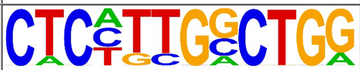 | 1e-8 | -2.009e+01 | 5.03% | 0.27% | 42.9bp (67.3bp) | Smad3(MAD)/NPC-Smad3-ChIP-Seq(GSE36673)/Homer(0.666) More Information \| Similar Motifs Found | motif file (matrix) |
| 10 * | 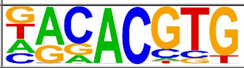 | 1e-5 | -1.338e+01 | 12.85% | 4.12% | 40.4bp (59.8bp) | Npas2/MA0626.1/Jaspar(0.943) More Information \| Similar Motifs Found | motif file (matrix) |
| 11 * | 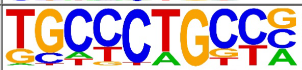 | 1e-4 | -1.070e+01 | 13.41% | 5.21% | 55.4bp (64.5bp) | ZNF416(Zf)/HEK293-ZNF416.GFP-ChIP-Seq(GSE58341)/Homer(0.757) More Information \| Similar Motifs Found | motif file (matrix) |
| 12 * | 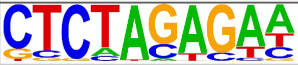 | 1e-3 | -8.670e+00 | 6.70% | 1.89% | 52.6bp (58.4bp) | Smad4/MA1153.1/Jaspar(0.701) More Information \| Similar Motifs Found | motif file (matrix) |
| 13 * | 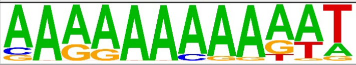 | 1e-3 | -7.176e+00 | 3.91% | 0.82% | 39.9bp (62.6bp) | ZNF384/MA1125.1/Jaspar(0.861) More Information \| Similar Motifs Found | motif file (matrix) |

**Supplementary Figure 4. Full list of de novo motif analysis shown in Figure 5g.** Full list of *de novo* motif analysis for the 190 downregulated TSS region DAR overlapped with DEG in Glis3$^{KO}$ *vs* WT by HOMER.

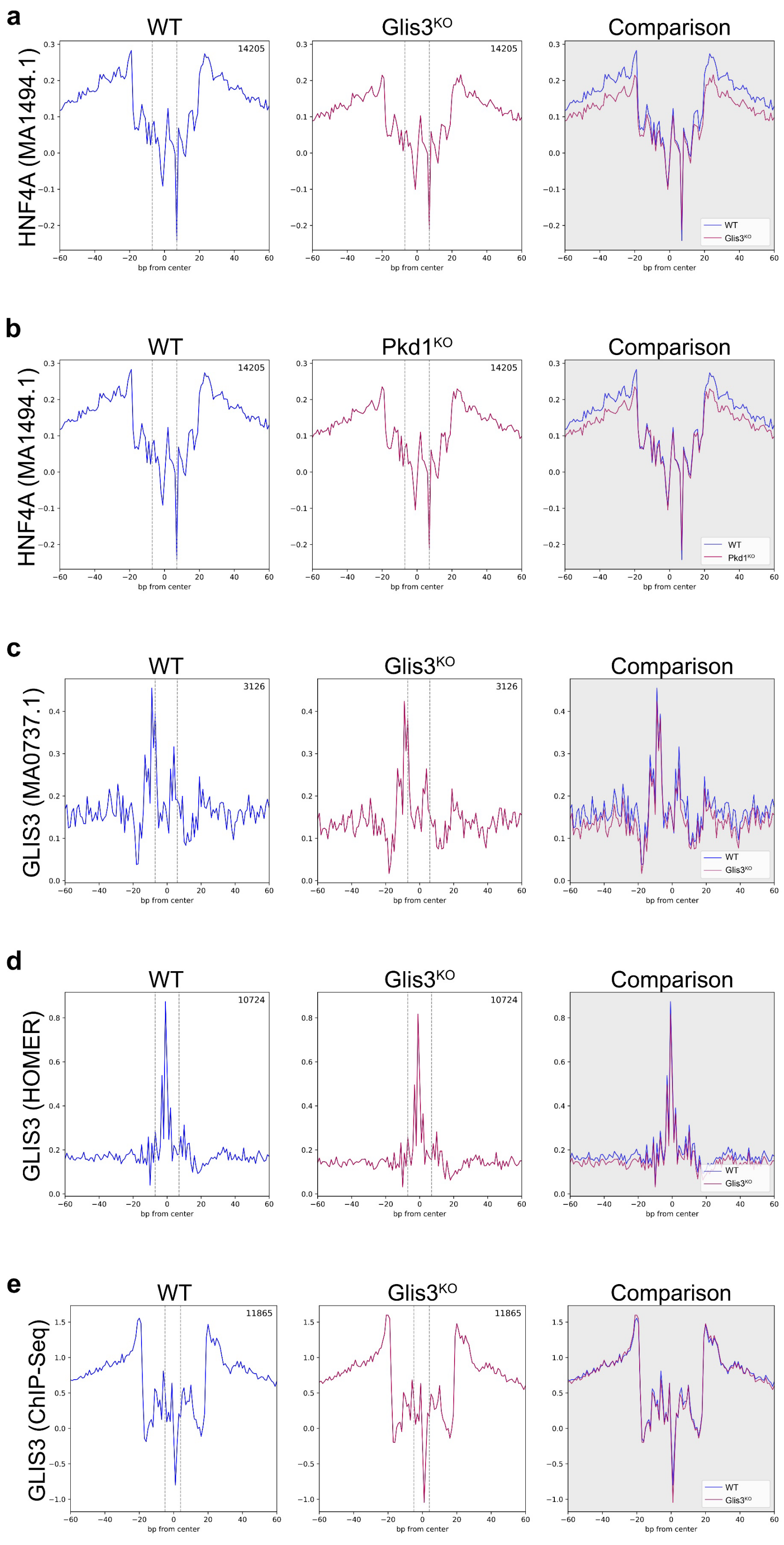


**Supplementary Figure 5. Footprinting analyses of Glis3 binding motifs.**
**a-b**, Footprints of a second binding motif for transcription factor Hnf4a (MA1494.1) in Glis3$^{KO}$ *vs* WT (**a**) and Pkd1$^{KO}$ *vs* WT (**b**). **c-e**, Footprints of different Glis3 binding motifs in Glis3$^{KO}$ *vs* WT. Sources of Glis3 binding motif from JASPAR (MA0737.1; **c**), HOMER (**d**), and published Glis3 ChIP-Seq (**e**).

| Gene Symbol | Description |
|---|---|
| Bhlhe40 | basic helix-loop-helix family, member e40 |
| P2ry2 | purinergic receptor P2Y, G-protein coupled 2 |
| Phldb1 | pleckstrin homology like domain, family B, member 1 |
| Lama1 | laminin, alpha 1 |
| Cyp2s1 | cytochrome P450, family 2, subfamily s, polypeptide 1 |
| Hpn | hepsin |
| Dab2ip | disabled 2 interacting protein |
| Ahdc1 | AT hook, DNA binding motif, containing 1 |
| Rhbdf1 | rhomboid 5 homolog 1 |
| Slc16a14 | solute carrier family 16 (monocarboxylic acid transporters), member 14 |
| Baiap2l2 | BAI1-associated protein 2-like 2 |
| Stk32a | serine/threonine kinase 32A |
| Lonrf1 | LON peptidase N-terminal domain and ring finger 1 |
| Rorc | RAR-related orphan receptor gamma |
| Ces1e | carboxylesterase 1E |
| Dbp | D site albumin promoter binding protein |
| Stk35 | serine/threonine kinase 35 |
| Fgfr4 | fibroblast growth factor receptor 4 |
| AI854703 | expressed sequence AI854703 |
| Pipox | pipecolic acid oxidase |
| Oplah | 5-oxoprolinase (ATP-hydrolysing) |
| Ddo | D-aspartate oxidase |
| Syne3 | spectrin repeat containing, nuclear envelope family member 3 |

**Supplementary Figure 6. Potential transcription targets of Glis3.**
List of 23 genes that are transcriptionally downregulated and have less open chromatin in the promoter region in Glis3$^{KO}$ *vs* WT and whose promoter directly bind to Glis3 based on published ChIP-Seq data.
